# Generalized strategy for engineering mammalian cell-compatible RNA-based biosensors from random sequence libraries

**DOI:** 10.1101/2022.10.18.512695

**Authors:** Everett R. Allchin, Jonah C. Rosch, Hyosung Kim, Ethan S. Lippmann

## Abstract

Fluorescent RNA-based biosensors are useful tools for real-time detection of molecules in living cells. These biosensors typically consist of a chromophore-binding aptamer and a target-binding aptamer, whereby the chromophore-binding aptamer is destabilized until a target is captured, which causes a conformational change to permit chromophore binding and an increase in fluorescence. The target-binding region is typically fabricated using known riboswitch motifs, which are already known to have target specificity and undergo structural changes upon binding. However, known riboswitches only exist for a limited number of molecules, significantly constraining biosensor design. To overcome this challenge, we designed a framework for producing mammalian cell-compatible biosensors using aptamers selected from a large random library by capture-SELEX. As a proof-of-concept, we generated and characterized a fluorescent RNA biosensor against L-dopa, the precursor of several neurotransmitters. Overall, we suggest that this approach will have utility for generating RNA biosensors that can reliably detect custom targets in mammalian cells.

## Introduction

Fluorescent biosensors convert a biological response into a measurable output, and when encoded into living cells, they can be used to monitor and measure molecular interactions, kinetics, and abundances.^1–4^ Historically, in-cell biosensing has been predominantly accomplished using proteins. Fluorescent proteins such as green fluorescent protein (GFP) engineered with recognition domains, and fluorescently-tagged proteins for proximity assays such as Förster resonance energy transfer (FRET), are frequently used as biosensors in research applications.^5–8^ While these approaches are incredibly useful, they have some distinct limitations. Fluorescent proteins are difficult to conjugate to non-protein molecules and can even prevent the function of some proteins after tagging onto the N- or C-terminus.^9,10^ When used as a reporter, the output loses temporal resolution as maturation of fluorescent proteins can take hours and even the quickest folding proteins can take 10 or more minutes.^11–13^ Fluorescent proteins also have long half-lives, which can further impact temporal resolution.^14^ Split GFP and GFP-based binders (GFAb) approaches seek to make GFP a sensor on its own with some success, but have many of the same limitations as fluorescent proteins in general.^15,16^ Temporal resolution is still restricted by protein folding and chromophore maturation as well as the near-irreversible nature of the complementation due to low off rates.^17–19^ Split GFP can also self-assemble leading to false positives.^19^ Unlike fluorescent proteins, FRET can detect small molecules and remains directly related to the number of targets. However, while this technique is very sensitive, the placement of the protein domains is relatively difficult and does not lend itself to facile detection of new targets.^20^

In contrast to fluorescent proteins, aptamer-based biosensors are fabricated using RNA oligonucleotides capable of binding to specific molecules.^21^ Aptamers have previously been converted into biosensors by coupling with a reporter gene. While initially unstructured, stabilizing interactions upon binding a ligand causes the aptamer to block binding of the ribosome or other translation elements, which reduces expression of a reporter gene. This method has been used in both bacteria and eukaryotes for biosensing.^22,23^ Efforts to use fluorescent aptamers for direct small molecule sensing were enabled by the development of Spinach, an RNA aptamer mimic of GFP,^24^ which has been used for diverse applications across RNA imaging^25,26^, small molecule sensing^27–29^, and real time transcription monitoring *in vitro* and *in vivo*.^30,31^ The original Spinach aptamer was converted to a fluorescent small molecule biosensor by inserting naturally-occurring riboswitch motifs into the parent fluorescent RNA structure. The insertion of the riboswitch motif destabilizes the aptamer to diminish chromophore binding, whereupon binding of the riboswitch motif to a target model induces a conformational change to restore chromophore binding, thus creating a “light up” construct that is activated by the riboswitch target. Since the development of the original Spinach aptamer, numerous advancements have been made in this area. For example, Spinach was generated using standard Systematic Evolution of Ligands by Exponential Enrichment (SELEX) approaches to isolate aptamers that could bind the DFHBI chromophore.^24^ A revisit to the SELEX process was used to select newer aptamers after expressing an enriched library of RNAs within *E. Coli* cells and using fluorescence-activated cell sorting (FACS) to isolate clones with high levels of fluorescence.^32^ This method was used to identify Broccoli, a markedly improved aptamer with nearly twice the brightness of Spinach and enhanced thermostability. Spinach requires higher Mg^2+^ concentrations than normally found in cells in order to fold properly, but Broccoli does not have this issue, which makes it more compatible for *in vivo* imaging.

The development of Broccoli revealed a disadvantage of these aptamers for use in mammalian cells. Aptamers in general exhibit suboptimal functions in cells due to RNA degradation and poor folding.^32^ Folding may be hindered by competing folding pathways, adjacent RNA sequences, thermal instability, or dependence on ion concentrations not present in cells.^33–35^ Faster degradation and lower expression levels can also lead to reduced aptamer concentrations in mammalian cells as compared to bacteria.^36^ This issue has been recently addressed using a new expression system called Tornado (Twister-ribozyme RNA for durable overexpression) that produces circularized aptamers to prevent common degradation pathways.^37^ RNA circularization can be achieved using a number of other methods including chemical and enzymatic methods (*in vitro* only) and both hairpin and cyclase ribozyme methods such as permuted introns and exons (PIE method).^38–40^ These modifications have improved stability of aptamers in mammalian cells and enabled further enhancements to biosensor performance.

However, despite these advances, a persistent limitation still exists with the small molecule recognition motifs. Specifically, recognition motifs to this point have been predominantly derived from naturally-occurring riboswitches. This limits detection solely to small molecules that have cognate riboswitch pairs and completely prevents the use of aptamer biosensors for non-native small molecules like synthetic drugs. Previous attempts to overcome this issue have randomized regions of naturally-occurring riboswitches to create riboswitch-like libraries for SELEX selection.^41^ This approach successfully produced biosensors in *E. coli* by splicing these SELEX hits into fluorogenic aptamer motifs such as Broccoli. However, to conserve the structure of the riboswitch scaffold, these libraries were limited to 23 random nucleotides, which lowers library diversity below what is typically recommended for deep selection.^42^ Hence, it is currently unknown if riboswitch-like aptamers selected from large libraries can function in fluorogenic RNA scaffolds to create real time biosensors.

Here, we sought to address this question by developing a new workflow that integrates aptamers selected from a capture SELEX workflow into a fluorogenic RNA scaffold (Figure 1). Capture SELEX is a technique that selects for conformational changing aptamers, making it ideal for identifying aptamers with biosensing characteristics. We focused on l-3,4-dihydroxyphenylalanine (L-dopa), a normally occurring amino acid precursor for neurotransmitters; a L-dopa biosensor was created from the previously mentioned riboswitch-like library, allowing us to make comparisons to our approach.^41^ We used capture SELEX to generate candidate RNA aptamers that could prospectively bind L-dopa and inserted a subset of these candidates into the Broccoli construct, followed by screening for fluorescent responses in cells. These tests led to the identification of aptamer 3F, which exhibited modest biosensing activity upon transient transfection. We then created a stable 3F-expressing Caco-2 cell line and carried out detailed characterizations of L-dopa biosensing dynamics, which showed L-dopa-dependent fluorescence increases using flow cytometry and real-time imaging. We further showed that inhibition of L-dopa metabolism with the small molecule carbidopa could prolong fluorescent signal. Last, we determined that the 3F biosensor had similar performance to the top L-dopa-sensing aptamer selected from the prior riboswitch-like library.^41^ Overall, our results establish a generalizable strategy for creating mammalian cell-compatible RNA biosensors against custom small molecule targets from fully random libraries.

**Figure 1:**
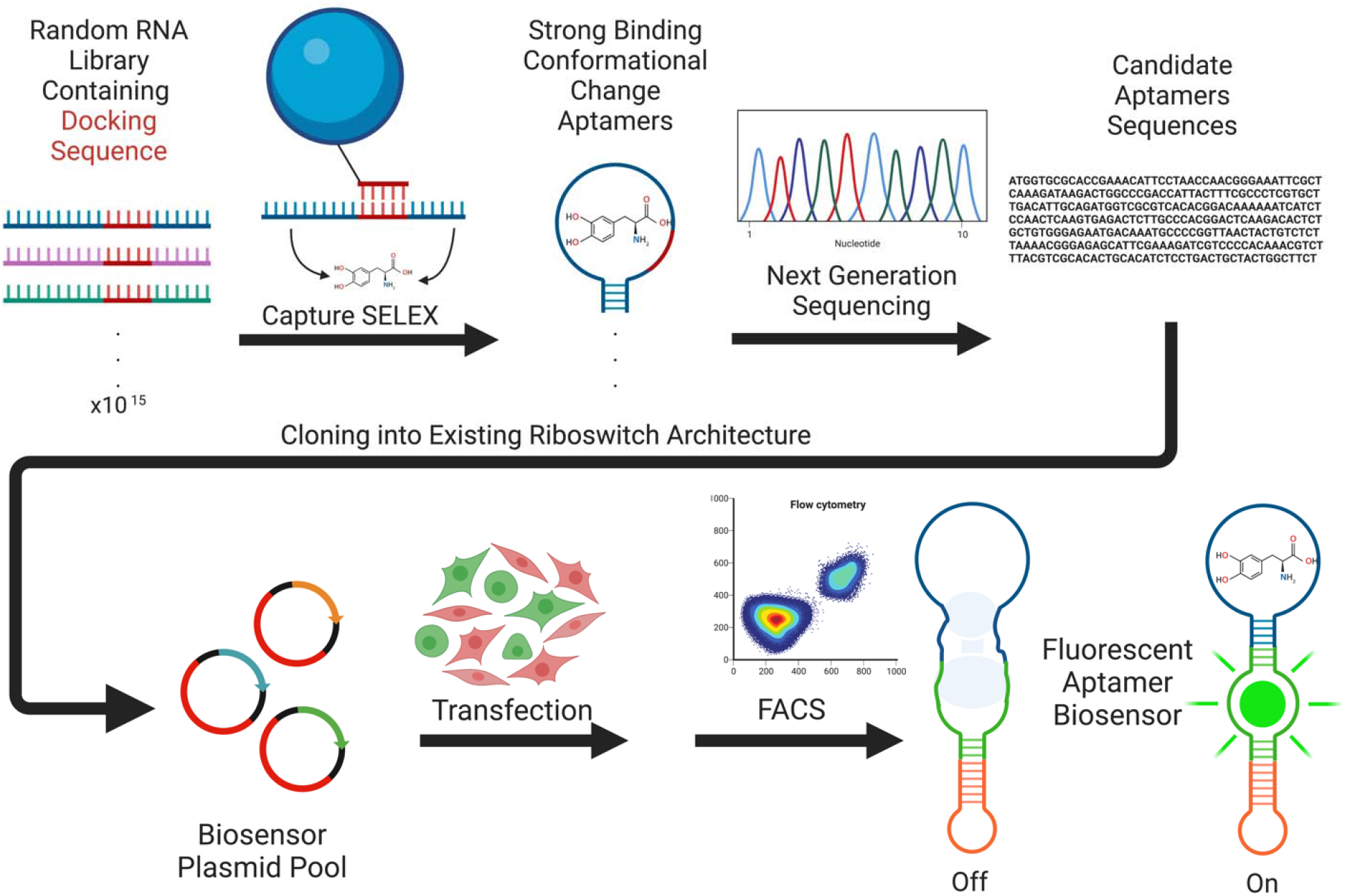
Overall workflow for biosensor fabrication. The schematic depicts the sequential steps of capture-SELEX, next-generation sequencing to identify candidate aptamers, and the assessment of these aptamers as L-dopa biosensors in a fluorescent construct. Schematic created with BioRender.

## Methods

### Nucleic acid sequences

All relevant sequences for the starting RNA library, capture oligonucleotide, primers, and gene blocks are found in Table S1.

### Capture SELEX

A DNA library was purchased from IDT with a 40 nucleotide random sequence and 10 nucleotide random sequence flanking a docking sequence and additional bases on either side as handles for primers T7_fwd and T7_rev, similar to a previously used library.^43^ Nucleotides were hand mixed to ensure equal incorporation of each base into the library. This library was PCR amplified to add a T7 promoter sequence using Taq polymerase and primers T7_fwd and T7_rev (40 cycles, denaturation: 95°C for 30 seconds, annealing: 60°C for 30 seconds, extension: 68°C for 30 seconds). PCR products were then *in vitro* transcribed with T7 polymerase (Lucigen). Capture SELEX was then carried out as follows. In the first round of selection, 10 nmol of RNA (∼1×10^15^ molecules) was annealed to the biotinylated capture oligo (IDT) at 65°C for 5 min before cooling to room temperature. This mixture was then incubated with 1 mL of 1mg/mL streptavidin-functionalized magnetic beads (M-270 Dynabeads). The beads were washed three times with SELEX buffer (40 mM HEPES, 350 mM KCl, 20 mM NaCl, 5 mM MgCl_2_) by agitating at 150 rpm for 5 minutes on a shaker plate. Beads were then washed with 1 mM L-dopa (Tocris bioscience) agitating at 150 rpm for 5 minutes to elute L-dopa binding aptamers. A magnet was used to concentrate the beads to one side and effluent was pipetted out. Eluted aptamers were precipitated with ethanol and reverse transcribed using a SunScript reverse transcription kit (Expedeon) according to the manufacturer’s instructions. 10 μL of reverse transcription products were then PCR amplified using Phusion polymerase (NEB) and T7_fwd and T7_rev (11 cycles, denaturing: 98°C for 10 seconds, annealing: 60°C for 15 seconds, extension: 72°C for 30 seconds). The PCR products were resolved on a 3% agarose gel, and the band with the appropriate size was extracted and *in vitro* transcribed to begin the next round of SELEX. For rounds 2-8, the volume of streptavidin-coated beads was reduced to 150 μL (0.15 mg of beads). For rounds 5-8, the concentration of L-dopa was reduced to 0.1 mM and the PCR amplification after reverse transcription of recovered sequences was increased to 22 cycles. For rounds 7-8, 0.1 mM tyrosine (Sigma) was included in the SELEX buffer for the washing step prior to incubation with L-dopa. After 8 total rounds, DNA from rounds 6 and 8 were submitted to the Vanderbilt Technologies for Advanced Genomics core facility for next generation sequencing on an Illumina NovaSeq6000. 12 aptamers were chosen from the sequencing results based on metrics described in the Results section. Analyses were performed using AptaSUITE software.^44^

### Biosensor pool cloning

A TORNADO Broccoli sequence expressed via a U6 promoter was inserted into a red fluorescent protein (RFP) backbone via Gibson assembly. The V21 CS2+ RFPbb plasmid (gift from Randall Moon; Addgene plasmid 17112) was PCR amplified with Phusion polymerase using primers RFPbb_fwd and FRPbb_rev (35 cycles, denaturing: 98°C for 10 seconds, annealing: 65°C for 10 seconds, extension: 72°C for 150 seconds). The pAV-U6+27-Tornado-Broccoli plasmid (gift from Samie Jaffrey; Addgene plasmid 124360) was PCR amplified with Phusion polymerase using primers TBins_fwd and TBins_rev (35 cycles, denaturing: 98°C for 10 seconds, annealing: 64°C for 10 seconds, extension: 72°C for 30 seconds). Gibson assembly was performed with these products using the Gibson assembly master mix (NEB E2611) with an incubation time of 20 minutes room temperature. Assembled plasmids were transformed into TOP10 competent cells (Invitrogen) by adding 5 μL Gibson reaction to comp cells on ice for 30 minutes. Bacteria were then heat shocked at 42°C for 30 seconds. 250 μL SOC (Corning) was then added and the cells were shaken at 250 rpm at 37°C for 1 hour. Cells were then plated on ampicillin plates and incubated at 37°C overnight. Colonies were picked for overnight culture in 3 mL of LB broth (KD medical) containing 50 μg/mL ampicillin (Mediatech/Cellgro – Corning MT61238RH). Plasmid DNA was recovered using a miniprep kit (Qiagen) according to the manufacturer’s instructions. This plasmid is referred to as RFP+TORNADO Broccoli.

Sequences were subsequently cloned into RFP+TORNADO Broccoli plasmid. Gene blocks for each of the 12 candidate aptamers were purchased from IDT. Each gene block contained the following from 5’ to 3’: Broccoli aptamer, candidate aptamer, Broccoli aptamer flanked by sites for the restriction enzymes NotI and SacII (Figure S1). 50 μL PCR reactions for each gene block were carried out with Phusion polymerase (NEB) using primers gblock_fwd and gblock_rev for 35 cycles (denaturing: 98°C 10s, annealing: 67°C 10s, extension: 72°C 30s). Each amplified gene block, as well as the Gibson assembled RFP+TORNADO Broccoli plasmid, was digested with NotI-HF and SacII enzymes (NEB) for 30 minutes at 37°C. Antarctic phosphatase (NEB) was added after 15 minutes to the RFP+TORNADO Broccoli digest only. 40 μL of each digest product were then ran on a 3% agarose gel and the appropriate band was gel extracted using a kit (NEB) according to the manufacturer’s instructions. Each gel extracted insert (one from each gene block) was then ligated to the gel extracted backbone from the RFP+TORNADO Broccoli digest in a separate 20 μL T7 ligation reaction (Thermo Fisher Scientific) at room temperature for 30 minutes (12 total reactions). 5 μL of ligation products were then transformed into competent cells (NEB). Transformation and plasmid DNA recovery were performed as previously described. Sequences were confirmed by Sanger sequencing picked colonies.

### HEK293 cell culture and initial assessment of candidate biosensors

HEK293 cells were cultured in DMEM (4.5 g/L glucose and sodium pyruvate) containing 6 mM L-glutamine, 10% FBS, 100 U/mL Penicillin, and 100 μg/mL streptomycin. 3×10^5^ cells were seeded into 12-well plates that had been coated with 1 mg/mL poly-L-ornithine (Sigma) for 60 minutes. At 90% confluency (after approximately 2 days), cells were transfected in opti-mem with Lipofectamine 3000 (Thermo Scientific) containing 2 μg of plasmid DNA. Cells remained in opti-mem overnight before being changed back to HEK medium. 48 hours post-transfection, cells were treated with 500 μM DFHBI (Tocris Bioscience) and 40 μM L-dopa for 30 minutes. Cells were then detached from the plate using TrypLE Express (Fisher Scientific). Cells were centrifuged at 300 rpm for 5 minutes, resuspended in FACS buffer (2% FBS in PBS), and kept on ice. Flow cytometry was performed on an Amnis Cellstream flow cytometer (Luminex) and analyzed with the accompanying Cellstream software. Cells were gated for single cells and 10,000 such events were recorded and analyzed.

### Generation of lentivirus and transduction of Caco-2 cells

Expression plasmid pLV[Exp]-U6>SAM Tornado Broccoli-CMV>mCherry, which integrates the TORNADO Broccoli construct with the 3F biosensor sequence, was synthesized by VectorBuilder. The transfer vector was incorporated into lentivirus using the lentiviral packaging plasmids (pMD2.G and psPAX2), which were a gift from Didier Trono (Addgene plasmids 12259 and 12260). HEK293T cells were plated at 60-70% confluence onto 6-well plates and cultured in DMEM supplemented with 10% fetal bovine serum (FBS). After overnight culture, the cells were co-transfected with 150 ng pMD2.G, 400 ng psPAX2, and 500 ng TORNADO Broccoli expression plasmid using Lipofectamine 3000 (Invitrogen). After 48 and 72 hours, the spent medium was collected and filtered through a 0.45 μm cellulose acetate filter before concentrating the supernatant using Lenti-X concentrator (TaKaRa). Caco-2 cells were cultured at <50% confluence and transduced with lentiviral particles. Selection was performed with FACS, with gating for live cells (as determined by DAPI exclusion) and high red fluorescence (see Figure S2 for gating strategy). Caco-2 cells stably expressing the 3F biosensor were maintained in DMEM (high glucose, sodium pyruvate, and L-glutamine) containing 10% FBS, 100 U/mL Penicillin, 100 μg/mL streptomycin, and 10x non-essential amino acids.

### Flow cytometry assessment of 3F biosensor performance in Caco-2 cells

12-well plates were pre-coated with 0.1% gelatin for 30 minutes. 3F-expressing Caco-2 were seeded into these plates at a density of 3×10^5^ cells/well. At 80% confluence, cells were treated in opti-mem with combinations of 40 μM DFHBI, 500 μM carbidopa (Spectrum Chemical Mfg.Corp.), and 500 μM L-dopa at varying time points. Treatment was continuous for the duration of the experiment(up to 1 hour for the maximum time point). All cells were then simultaneously collected from the plate at the endpoint using TyrpLE Express. Cells were centrifuged at 300 rpm for 5 minutes, resuspended in FACS buffer (2% FBS in PBS), and kept on ice. Flow cytometry was performed on an Amnis Cellstream flow cytometer and analyzed with the accompanying Cellstream software. Cells were gated for single cells and 10,000 such events were recorded and analyzed.

### Assessment of 3F biosensor performance in Caco-2 cells using fluorescence microscopy

12-well plates were pre-coated with 0.1% gelatin for 30 minutes. 3F-expressing Caco-2 cells were seeded into these plates at a density of 3×10^5^ cells/well. At 80% confluence, cells were treated in opti-mem with 40 μM DFHBI, 500 μM carbidopa, and/or 500 μM L-dopa continuously for 2 hours and imaged using a Leica DMi8 fluorescence microscope. Identical acquisition settings were used for all conditions. Images were analyzed in ImageJ.

### DGR aptamer cloning, transfection, and evaluation

The DGR-II CM-5 sequence^41^ was cloned into the RFP+TORNADO Broccoli plasmid. The DGR-II CM-5 gene block was purchased from IDT and contained the aptamer sequence coupled with Broccoli and flanked by sites for the restriction enzymes NotI and SacII (Table S1). A 50 μL PCR reaction for the gene block was carried out with Phusion polymerase (NEB) using primers DGR_gblock_fwd and DGR_gblock_rev for 35 cycles (denaturing: 98°C 10s, annealing: 67°C 10s, extension: 72°C 30s). The amplified gene block, as well as the Gibson assembled RFP+TORNADO Broccoli plasmid, was digested with NotI-HF and SacII enzymes (NEB) for 30 minutes at 37°C. Antarctic phosphatase (NEB) was added after 15 minutes to the RFP+TORNADO Broccoli digest only. 40 μL of digest product was then ran on a 3% agarose gel and the appropriate band was gel extracted using a kit (NEB) according to the manufacturer’s instructions. The gel extracted insert was then ligated to the gel extracted backbone from the RFP+TORNADO Broccoli digest in a 20 μL T7 ligation reaction (Thermo Fisher Scientific) at room temperature for 30 minutes. 5 μL of ligation product was then transformed into competent cells (NEB). Transformation and plasmid DNA recovery were performed as previously described. Sequences were confirmed by Sanger sequencing picked colonies.

To evaluate performance, Caco-2 cells were transfected with the DGR or 3F plasmid (1 μg DNA) using Lipofectamine 3000 in opti-mem as described earlier. Cells were maintained opti-mem overnight before being switched back to standard Caco-2 medium. 48 hours post-transfection, cells were treated with 500 μM DFHBI and 40 μM L-dopa for 20 minutes. Cells were then detached from the plate using TrypLE Express, centrifuged at 300 rpm for 5 minutes, resuspended in FACS buffer, and placed on ice. Flow cytometry was performed on an Amnis Cellstream flow cytometer Cells were gated for single cells and 10,000 events were recorded and analyzed. Cells treated solely with DFHBI were used as a negative control.

## Results and Discussion

### Capture-SELEX selection of candidate L-dopa-binding aptamers

The initial aptamer library was based on a previous design used for successful riboswitch selection in an RNA Capture-SELEX workflow^43^ and is inspired by another pool used for DNA Capture-SELEX.^45^ A 50 nucleotide random region was split into a 40 nucleotide and 10 nucleotide region by a 13 nucleotide docking sequence. The docking sequence was then annealed to its complement—the capture oligo—which is bound to a bead via a streptavidin-biotin interaction. Sequences were then eluted by washing with the target molecule. To elute, sequences must undergo a change in structure upon interacting with the target that effectively yields strand displacement of the docking sequence and the capture oligo. This structural change upon binding is intended to mimic riboswitch behavior. Capture SELEX was conducted on the RNA library for 8 rounds (see Table S2 for parameters). The first 4 rounds used 1 mM L-dopa to elute prospective aptamers. At round 5, L-dopa concentration was reduced to 0.1 mM to increase stringency by requiring higher affinity binding to L-dopa. At round 7, stringency was further increased by washing with tyrosine prior to elution with L-dopa. Tyrosine was chosen for this counterselection step due to its chemical similarity to L-dopa (differing by a single hydroxyl group). When round 8 of selection was completed, eluted nucleic acid pools from rounds 6 and 8 were subjected to high throughput sequencing. 12 candidate aptamers (Table 1) were chosen for further analysis based on three criteria: maximum enrichment between rounds 6 and 8, most abundant sequences in the 8^th^ round, or family analysis which groups aptamers that are similar in sequence and predicted structure. Only aptamers containing the docking sequence were chosen, since without the docking sequence, aptamers are likely unable to bind to the capture oligo and therefore were not selected for their ability to undergo a conformational change when binding L-dopa. These sequences may have arisen during PCR or other amplification/purification steps and were therefore excluded from consideration.

**Table 1.**
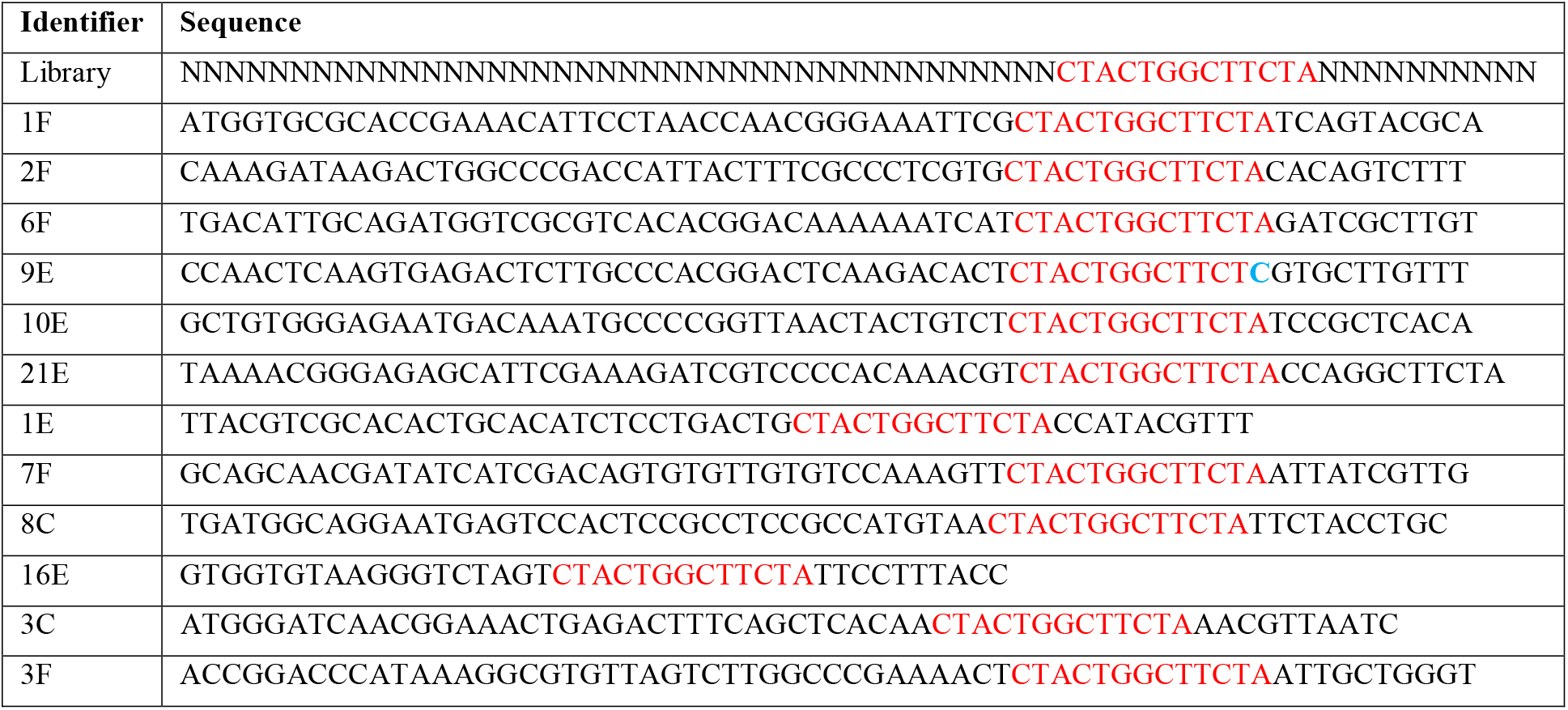
Candidate L-dopa aptamers identified by capture SELEX. Labels represent if the sequence was selected based on enrichment between rounds 6 and 8 (E), count in the 8^th^ round (C), or a family analysis that groups aptamers that are similar in sequence and predicted structure (F). Only aptamers containing the docking sequence were chosen (shown in red). A sequence that has a mutation in the docking sequence is shown in blue.

### Screening biosensor activity in mammalian cells

To convert the L-dopa-binding aptamers into biosensors, a design from a previous fluorescent aptamer biosensor based on the S-adenosylmethionine (SAM) riboswitch was used.^37^ This design uses a TORNADO construct to circularize the RNA structure, and a Broccoli fluorescent domain is present in the center of the RNA that is only capable of binding the chromophore DFHBI upon structure completion when the SAM aptamer binds its target. The 12 candidate L-dopa aptamers were individually cloned into this expression cassette in place of the SAM aptamer, and the entire RNA was expressed under control of the U6 promoter. Additionally, the expression cassette contains a separate CMV promoter region for constitutive expression of red fluorescent protein (RFP), which was used as a transfection control. Each of these potential biosensors was screened by transfection into HEK293 cells, followed by quantification of fluorescence using flow cytometry. From these initial tests, we chose three candidate aptamers (9E, 8C, and 3F; one from each category criteria mentioned above) that showed a promising increase in fluorescence with L-dopa. These candidates were subjected to re-evaluation with additional biological replicates, and of these, the 3F aptamer was selected as the best candidate based on a reproducible and statistically significant 1.2-fold change in fluorescence in the presence of L-dopa (Figure 2).

**Figure 2:**
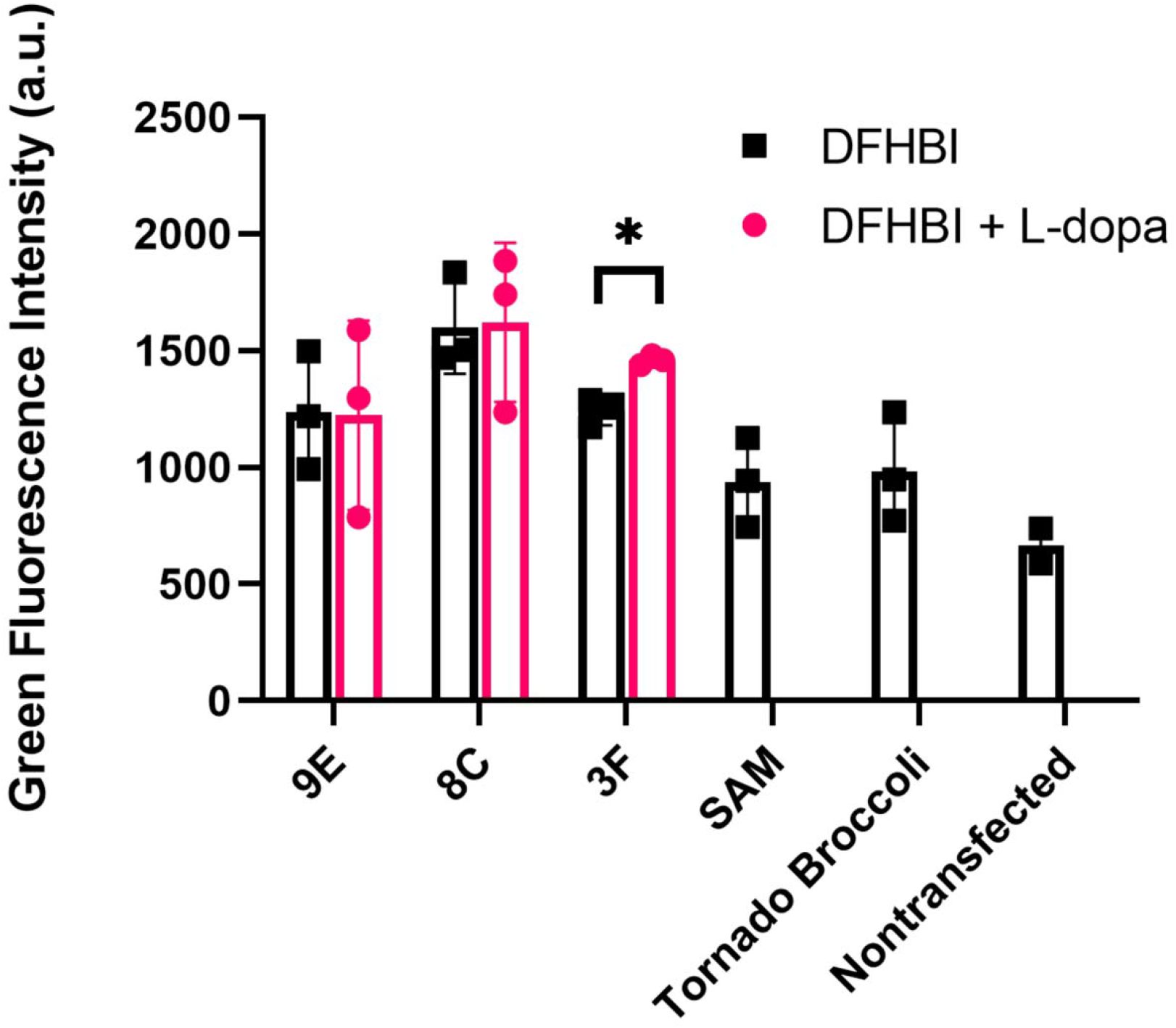
Evaluation of candidate L-dopa biosensors in HEK293 cells. Cells transfected with each prospective biosensor were incubated with DFHBI and L-dopa for 30 minutes, and fluorescence was measured using flow cytometry. Data are presented as mean ± standard deviation from N=3 biological replicates. Statistical significance was calculated using the student’s unpaired t-test (*, p<0.05). Untransfected cells, as well as cells transfected with plasmids encoding the SAM biosensor or the original Tornado Broccoli sequence, were assayed as controls in the presence of DFHBI alone.

### Characterization of biosensor candidate 3F

3F was next assessed in-depth as a potential L-dopa biosensor in Caco-2 cells. This cell line was chosen because, unlike HEK293 cells, Caco-2 cells are predicted to express the enzyme L-dopa decarboxylase that is responsible for metabolizing L-dopa into dopamine.^46^ This enzyme can be inhibited with the small molecule carbidopa, which provides an interesting context to study biosensor performance. To overcome limits of low transfection efficiency, we used lentiviral transduction to create a Caco-2 line with stable expression of the 3F biosensor. Then, we examined biosensor dynamics over time. We first treated the 3F-expressing Caco-2 cells with DFHBI and L-dopa and measured fluorescence at 10-minute intervals by flow cytometry (Figure 3A). Relative to cells treated with DFHBI alone, we found that fluorescence significantly increases and peaks at 10 and 20 minutes after simultaneous L-dopa and DFHBI addition, before dropping back to levels that were statistically insignificant compared to the negative control. To compare performance to the L-dopa aptamer (termed DGR) selected from a riboswitch-like library, we cloned DGR into the same Broccoli framework, transfected Caco-2 cells with DGR or 3F constructs, and measured fluorescence by flow cytometry at the ideal 20-minute time point (Figure 3B). Here, we did not observe any statistically significant differences between DGR or 3F, with both constructs yielding ∼1.2-fold increase in fluorescence in the presence of L-dopa and DFHBI versus DFHBI alone.

**Figure 3:**
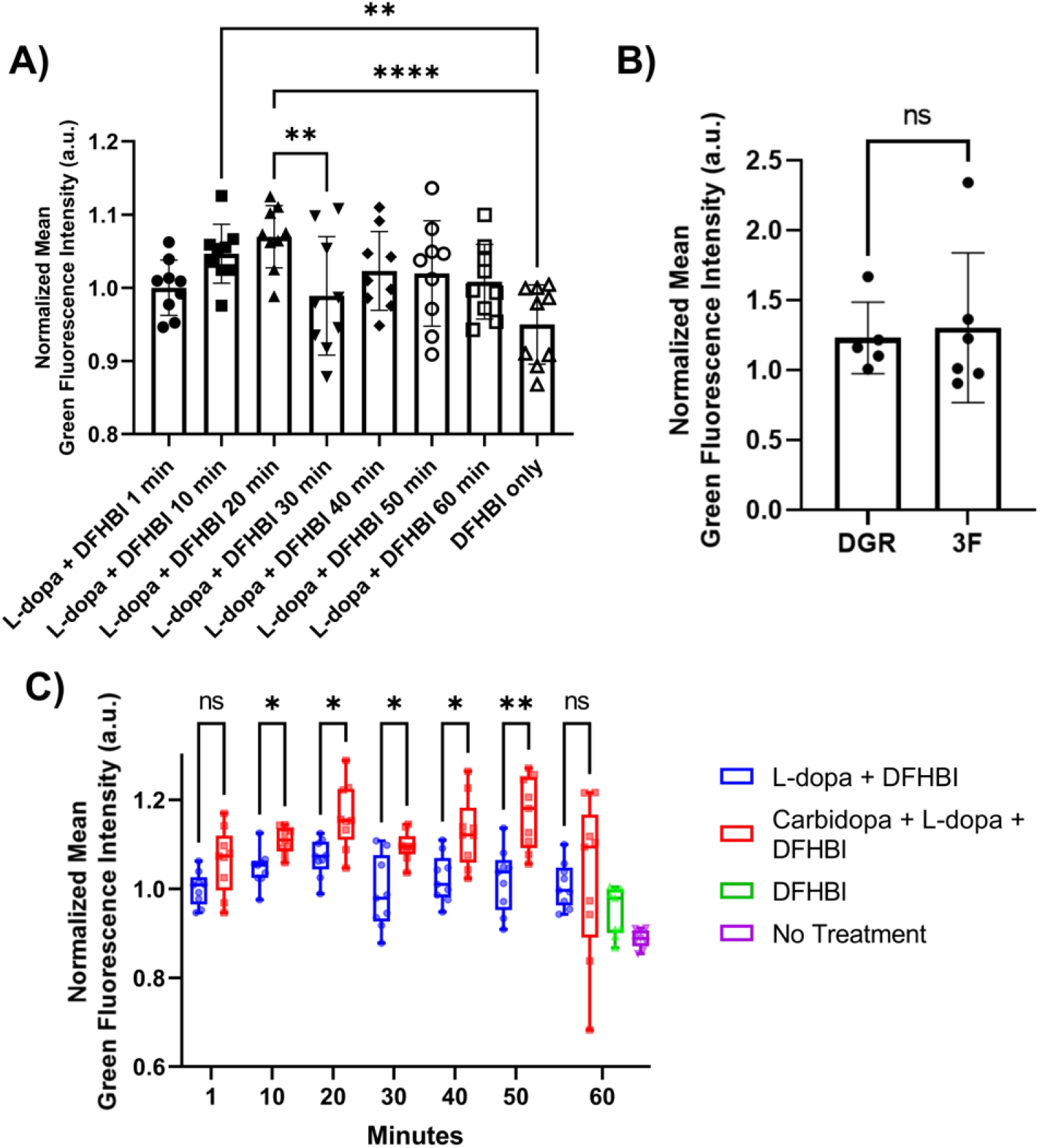
Evaluation of 3F biosensor in Caco-2 cells using flow cytometry. **A)** Cells stably expressing the 3F biosensor were incubated with DFHBI and L-dopa and then analyzed for fluorescence by flow cytometry at the indicated time points. For each sample, green fluorescence from the biosensor was first normalized to red fluorescence from RFP. These values were then further normalized to the 1-minute time point to assess temporal changes to L-dopa-induced fluorescence. Data are presented as mean ± standard deviation from N=9 biological replicates. Statistical significance was calculated using a one-way ANOVA with Sidak’s multiple comparisons at each time point (**, p<0.01; ****, p<0.0001). Cells that only received DFHBI served as a reference for fluorescence intensity. **B)** Cells transfected with DGR or 3F biosensor were incubated with DFHBI and L-dopa for 20 minutes, then analyzed by flow cytometry. Fold change was calculated relative to cells treated with DFHBI alone. Data are presented as mean ± standard deviation from 5-6 biological replicates. Statistical significance was calculated using a student’s unpaired t-test (ns, p>0.05). **C)** Under the same conditions and assessments as panel A, cells stably expressing the 3F biosensor were additionally treated with carbidopa. For each sample, green fluorescence from the biosensor was first normalized to red fluorescence from RFP. These values were then further normalized to DFHBI and L-dopa treatment at the 1-minute time point. Data are presented as mean ± standard deviation from N=9 biological replicates. Statistical significance was calculated using a two-way ANOVA (*, p<0.05; **, p<0.01). Untreated cells, as well as cells that only received DFHBI, served as a reference for fluorescence intensity.

Hypothesizing that the previously observed decrease in 3F biosensor fluorescence was due to L-dopa metabolism by dopa decarboxylase, we examined biosensor performance in the presence of carbidopa. Here, using pairwise comparisons, we determined that fluorescence was significantly increased by carbidopa treatment through 50 minutes (Figure 3C); at the 60-minute time point, differences were no longer significant due to increased heterogeneity in the carbidopa-treated samples. To confirm these outcomes, we additionally assessed fluorescence via microscopy in living cells. When imaging 3F-expressing Caco-2 cells treated with DFHBI alone or DFHBI and L-dopa, we could easily visualize increased green fluorescence in the L-dopa-treated cells. In general, fluorescence was uniformly induced throughout the cells at low levels, and brighter puncta were also observed in sporadic locations (Figure 4A), which is similar to the original SAM biosensor. As expected from the flow cytometry experiments, this fluorescence was lost over time, whereas the addition of carbidopa extended the duration of fluorescent signal (Figure 4B). Together, these data suggest that the 3F biosensor is specific for L-dopa and faithfully detects time-dependent L-dopa levels inside living cells.

**Figure 4:**
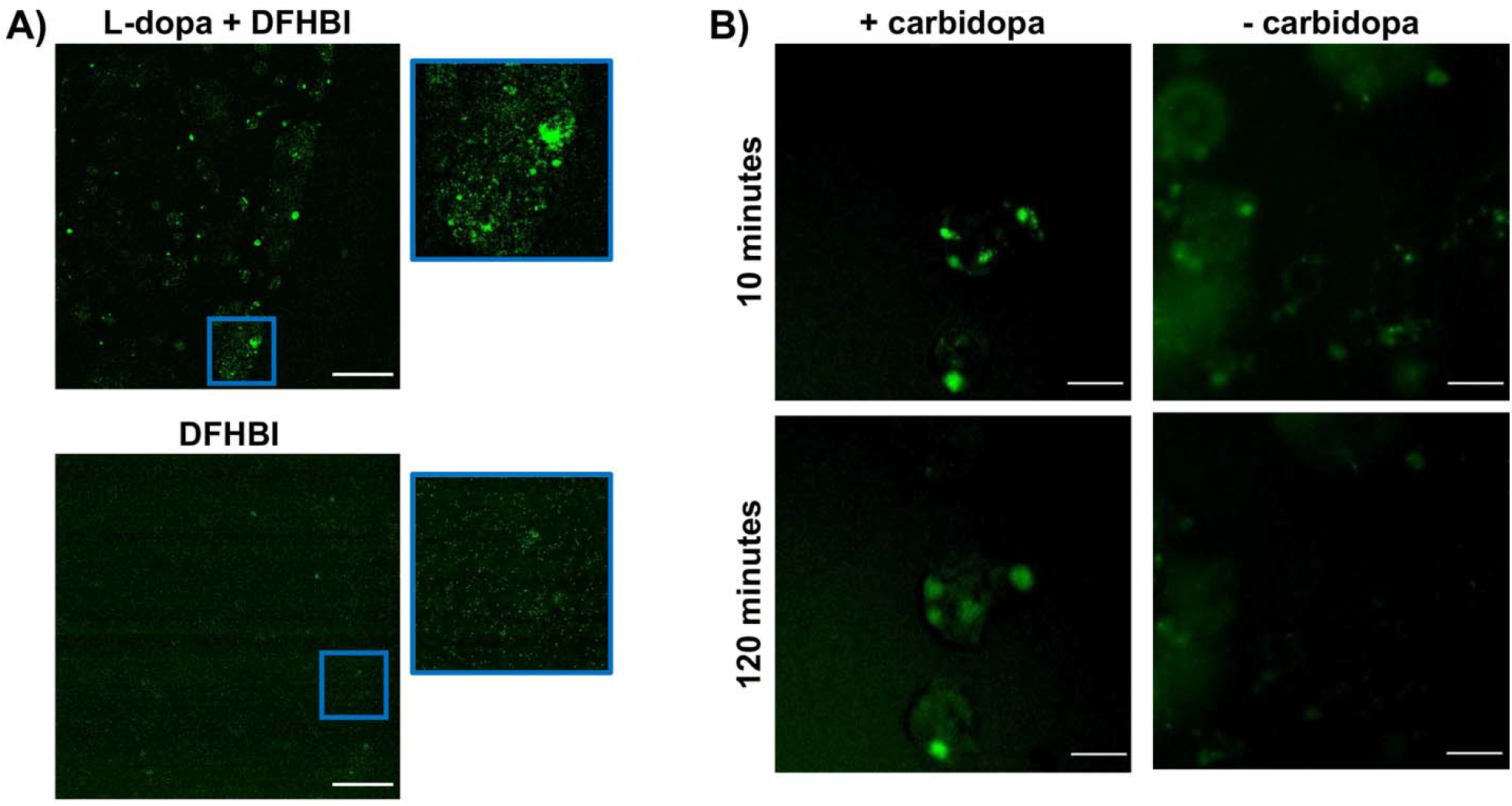
Evaluation of 3F biosensor in Caco-2 cells using fluorescence microscopy. **A)** Representative images of cells stably expressing the 3F biosensor and treated with DFHBI alone or DFHBI and L-dopa. Images were acquired after 10 minutes of treatment. An inset from each image highlights the specificity of the fluorescent signal to L-dopa treatment. Scale bars, 250 μm. **B)** Representative images at 10 minutes and 2 hours after treatment with DFHBI and L-dopa, with or without carbidopa. The same location was imaged for both samples. Scale bars, 50 μm.

## Conclusions

Our study shows that structure-switching aptamers selected from random libraries can be effectively integrated into existing RNA biosensor frameworks. Without any optimization, the 3F biosensor could produce a 1.2-fold change in fluorescence in the presence of L-dopa, which is comparable to Broccoli-based biosensors against other targets^47^ and the alternate L-dopa biosensor DGR (Figure 3B). We posit that further engineering of the ligation sequence between Broccoli and the L-dopa aptamer (which is currently a single A-U base pair, similar to the SAM biosensor^37^), use of higher performing chromophores (e.g. DFHBI-1T)^48^, and integration with alternate fluorescent aptamer sequences (such as red or orange Broccoli^49^, which may have reduced background in cells) could produce improvements in biosensor performance without requiring changes to the selection or screening process. Overall, we anticipate that this experimental framework will be useful for developing biosensors from random nucleic acid libraries.

## Author contributions

ERA, JCR, and ESL conceived the study. ERA carried out most experiments with initial assistance from JCR. HK provided support on lentiviral vector design and assisted with viral transductions. ESL supervised all work. ERA and ESL wrote the manuscript, and all authors read and approved its final format.

## Acknowledgments

This work was supported by a Ben Barres Early Career Acceleration Award from the Chan Zuckerberg Initiative (grant 2018-191850 to ESL). Support for the VANTAGE core facility was provided in part by a Clinical and Translational Science Award (5UL1 RR024975), the Vanderbilt Ingram Cancer Center (P30 CA68485), the Vanderbilt Vision Center (P30 EY08126), a CTSA award from the National Center for Advancing Translational Sciences (UL1 TR002243), and the National Center for Research Resources (G20 RR030956). Support for the Vanderbilt flow cytometry core was provided in part by the Vanderbilt Ingram Cancer Center (P30 CA68485) and the Vanderbilt Digestive Disease Research Center (P30 DK058404).

## Notes

### Competing Interest Statement

The authors have declared no competing interest.

