## Supplemental information for "Generalized strategy for engineering mammalian cell-compatible RNA-based biosensors from random sequence libraries"

**Table S1. Relevant nucleic acid sequences.**

| **Label** | **Sequence** |
| --- | --- |
| RFPbb_fwd primer | AAATAGGCCCTCTTCCTGCCCGACCTTGGAGCGAATTAAAAAACCTCCC |
| RFPbb_rev primer | TGTAGTTAATGATTAACCCGCCATGCTACTTCGCCAATGCATTGGGCCCGGTACCC |
| Tbins_fwd primer | TCCAAGGTCGGGCAGGAAGAGGGCCTATTTCCCATGATTCCTTCATA |
| Tbins_rev primer | TGTAGTTAATGATTAACCCGCCATGCTACTTATCTACGTA |
| T7_fwd primer | GTATAATACGACTCACTATAGGGCTTCTACCGCGGCCGC |
| T7_rev primer | ACACACGGACTTACGCCGCGG |
| Gblock_fwd primer | TAAAATGGGAGGGGGCGGG |
| Gblock_rev primer | GGCATTGGCAGTGTTCTACAGTCC |
| DGR_gblock_fwd | CCTAACCATGCCGAGTGCGGCCGCCGGAGACGGTC |
| DGR_gblock_rev | GCAGTGTTCTACAGTCCACGCCGACCGCGGCGGAGCCCACACTCT |
| Library | CTTCTACCGCGGCCGCNNNNNNNNNNNNNNNNNNNNNNNNNNNNNNNNNNNNNNNNCTACTGGCTTCTANNNNNNNNNNCCGCGGCGTAAGTCCGTGTGT |
| DGR gene block | GCGGCCGCCGGAGACGGTCGGGTCTATATGACTCCGTAATTCGCGTGGATATGGCACGCAGTAAGGTATTGGGCACCGTAAATGTCCATATGTATATATCGAGTAGAGTGTGGGCTCCGC |
| Capture oligo | TAGAAGCCAGTAG /3BioTEG/ |

**Table S2. Parameters used for capture-SELEX rounds.**

| **Round** | **Eluent** | **Counter-eluent** | **Washes** | **RNA input (µg)** |
| --- | --- | --- | --- | --- |
| 1 | 1 mM L-Dopa | - | 3 | 334 |
| 2 | 1 mM L-Dopa | - | 3 | 52.6 |
| 3 | 1 mM L-Dopa | - | 3 | 8.1 |
| 4 | 1 mM L-Dopa | - | 3 | 5.76 |
| 5 | 0.1 mM L-Dopa | - | 3 | 5.17 |
| 6 | 0.1 mM L-Dopa | - | 3 | 1.07 |
| 7 | 0.1 mM L-Dopa | 0.1 mM Tyrosine | 2 | 19.4 |
| 8 | 0.1 mM L-Dopa | 0.1 mM Tyrosine | 3 | 22.4 |

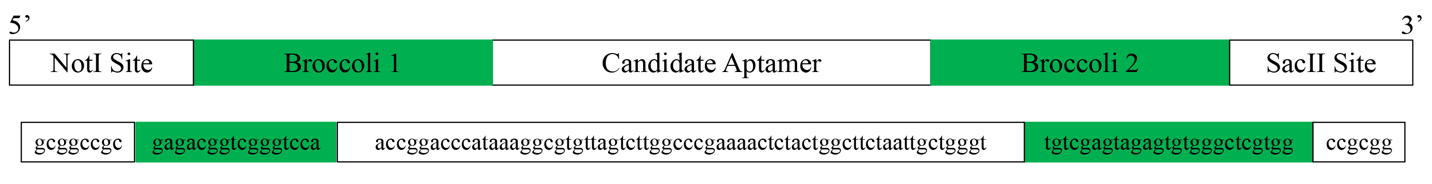

**Figure S1: Gene block design for inserting candidate aptamers into the Broccoli biosensor.** Sequence for the 3F aptamer is shown as an example.

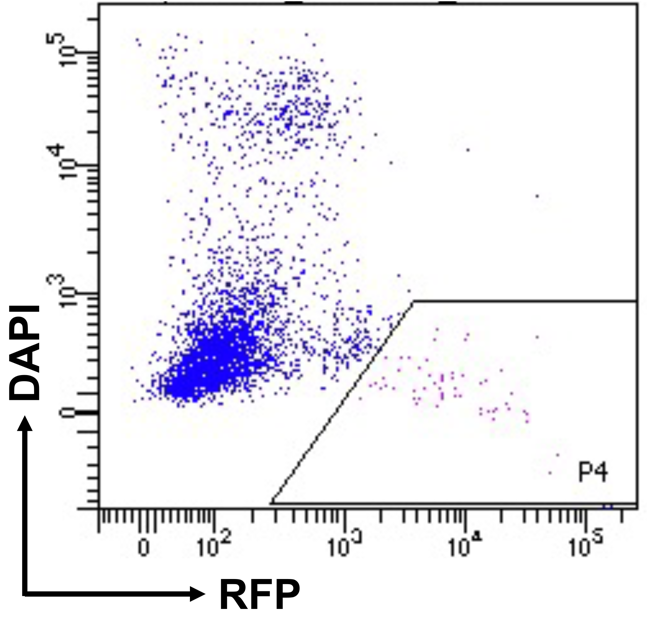

**Figure S2. Gating strategy for sorting 3F-expressing Caco-2 cells based on red fluorescent protein expression.** After compensation controls were applied, the P4 gate was used to select cells with elevated expression of RFP and exclusion of DAPI.

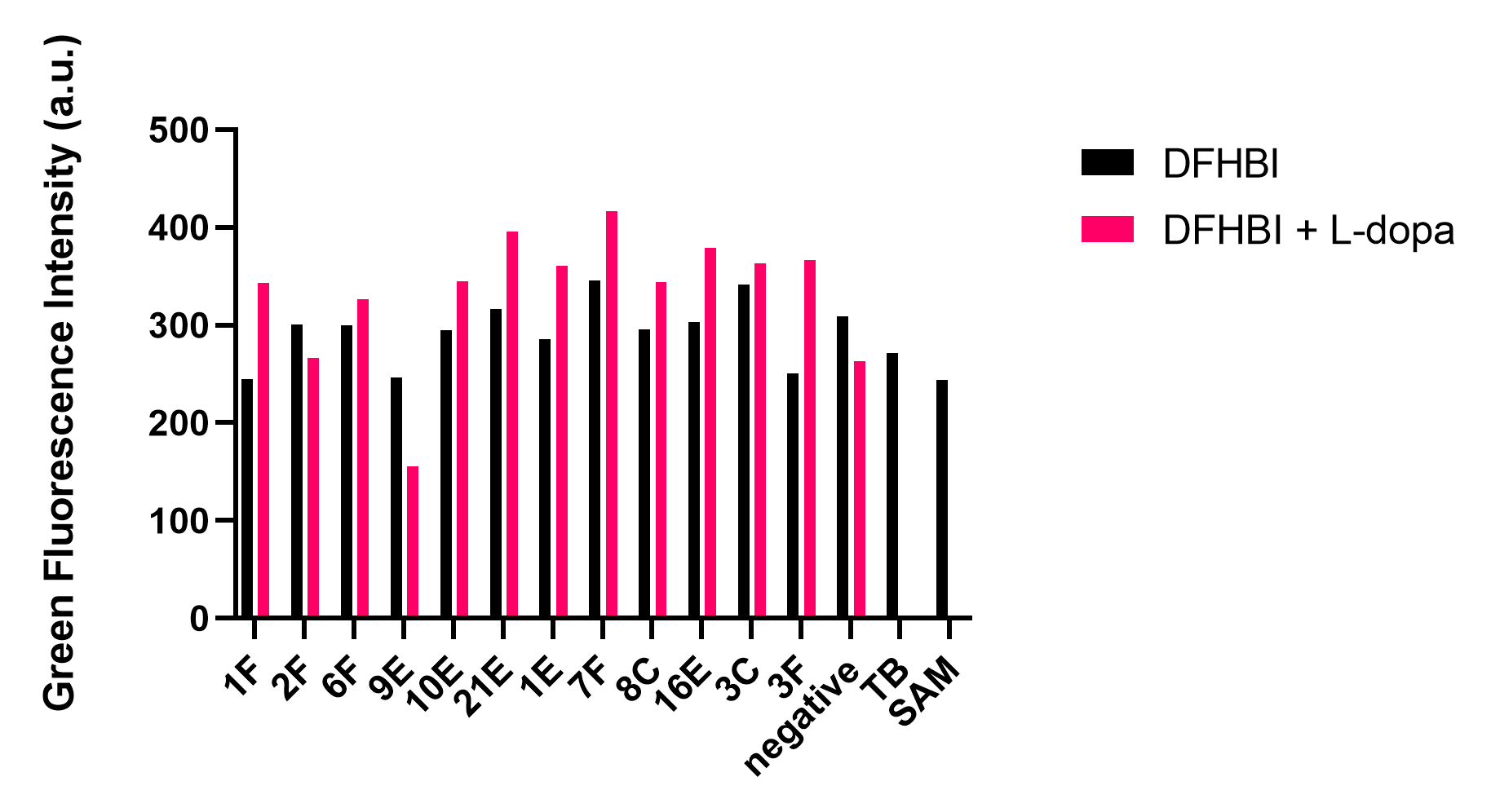

**Figure S3. Screening candidate aptamer biosensors after transfection into HEK293 cells.** Each sample represents a biological N=1 for each candidate, with and without L-dopa treatment. TB indicates the Tornado Broccoli construct without any inserted aptamer and SAM indicates the SAM aptamer in the Tornado Broccoli construct.
